# Integrated transcriptomic, metabolomic, and lipidomic analyses reveal a unique lipid profile of regulatory T cells upon activation

**DOI:** 10.1101/2024.12.15.627377

**Authors:** Yohei Sato, Misa Yura, Alana Chandler, Yamato Hanawa, Akihito Tsubota

## Abstract

Regulatory T cells (Tregs) exhibit stable FOXP3 expression and regulate the immune response through suppressive activity. Their unique metabolic properties include reduced glycolysis and increased oxidative phosphorylation. We combined transcriptomic, metabolomic, and lipidomic analyses to dissect the metabolic dynamics of Tregs upon activation. Combined metabolomic and lipidomic analyses showed that freshly isolated and activated Tregs had distinct metabolomic and lipidomic properties, respectively. Activated Tregs contained omega-3 long-chain polyunsaturated fatty acid (PUFA)-rich diglycerides and triglycerides. These were supported by transcriptomics data, showing upregulation of PPAR-alpha and PPAR-gamma. Activated Tregs exhibited increased ceramide production, consistent with the upregulation of ceramide synthase and sphingomyelin synthase. Confocal microscopy revealed that Tregs were enriched in lysosomes and peroxisomes compared with effector T cells upon activation. Our data confirm the unique metabolic properties of Tregs, especially those characterized by omega-3 long-chain PUFA-rich triglycerides and ceramides, together with enriched lysosomes and peroxisomes, which correspond to metabolic alterations.

## Introduction

Regulatory T cells (Tregs) control immune responses and are characterized by stable FOXP3 expression (Fontenot et al., 2003; Hori et al., 2003; Sakaguchi et al., 1995). Originally, FOXP3+ Tregs were identified in the CD4+CD25+CD127^low^ populations (Liu et al., 2006). Compared with effector T cells (Teffs), Tregs have unique metabolic properties, such as reduced glycolysis and increased oxidative phosphorylation (OXPHOS), which are supported by mitochondrial proliferation (Procaccini et al., 2016). To date, metabolic alterations in Tregs have primarily been analyzed with regard to carbohydrate and amino acid metabolism (Matias et al., 2021), whereas lipid metabolism has not been fully elucidated. T cells have been proposed to synthesize and utilize lipids for energy production and prolonged survival upon activation; however, the biological role of lipid metabolism in Tregs remains unknown (Pinzon Grimaldos et al., 2022). Previous studies have suggested that oleic acid can induce Treg cell differentiation (Lin et al., 2024; Pompura et al., 2021), but its lipidomic profile has not yet been identified. In the present study, integrated transcriptomic, metabolomic, and lipidomic analyses were performed to understand the unique lipidomic profile of Tregs. As PUFAs are metabolized in peroxisomes (Di Cara et al., 2023), we analyzed organelles, including peroxisomes, lysosomes, and mitochondria. Our findings demonstrate the importance of lipid metabolism in Treg function.

## Results

### Tregs had a distinct transcriptomic profile

Tregs and Teffs isolated from frozen peripheral blood mononuclear cells (PBMCs) were activated by T cell receptor (TCR) stimulation in the presence of recombinant human IL-2 (Figure S1A), as revealed by fluorescence-activated cell sorting (FACS). RNA and metabolites were isolated from multiple donors and pooled for further analysis (Figure S1B). The RNA-sequencing (RNA-seq) results are shown in Table S1. Differentially expressed genes (DEGs) are shown in Tables S2 (freshly isolated Tregs vs. Teffs) and S3 (activated Tregs vs. Teffs). Tregs had a distinct gene expression profile compared with Teffs, either before or after TCR stimulation (Figure S2A). While freshly isolated Tregs showed distinct gene expression, TCR stimulation drastically altered the gene expression profiles in both Teffs and Tregs (Figure S2B). As expected, freshly isolated Tregs had a distinct gene expression profile compared with freshly isolated Teffs (784 upregulated and 132 downregulated genes), mainly characterized by *FOXP3, IL2RA, CTLA-4, IKZF2,* and *LRRC32* expression (Figure S2C and Table S2). Moreover, activated Tregs had a distinct gene expression profile compared with activated Teffs (1282 upregulated and 351 downregulated genes), which included FOXP3-related genes, similar to freshly isolated Tregs (Figure S2D and Table S3). Pathway analysis identified significant enrichment of OXPHOS-related genes among the DEGs in activated Tregs (*p*<0.05) but not in freshly isolated Tregs. Interestingly, *RRARG* and *CD36* were upregulated in activated Tregs but not in freshly isolated Tregs. Transcriptomic analysis of activated Tregs strongly indicated that they may have unique metabolomic features, especially with regard to lipid metabolism, upon TCR stimulation. Therefore, we performed metabolomic and lipidomic analyses of freshly isolated and activated Tregs.

### Freshly isolated Tregs had a distinctive metabolomic profile

Next, we analyzed the metabolomic profile of Tregs, focusing on hydrophilic molecules, including carbohydrates and amino acids. The metabolomic profiles are shown in Tables S4 (relative quantification) and S5 (absolute quantification). A total of 203 (148 cations and 55 anions; Figures 1A and 1B) and 66 metabolites (43 cations and 23 anions; Figures S3A and S3B) were detected in the relative and absolute quantification, respectively. Differentially accumulated metabolites (DAMs) in freshly isolated and activated Tregs are presented in Tables S6 and S7, respectively. Freshly isolated Tregs had a distinctive metabolomic profile including 54 DAMs compared with Teffs (Figure 1C and Table S6). Activated Teffs and Tregs exhibited similar metabolomic profiles including 21 DAMs (Figure 1D and Table S7). As described previously, freshly isolated and activated Tregs exhibited a downregulation of the glycolytic pathway and upregulation of the TCA cycle (Figures S4A and S4B). Similarly, RNA-seq revealed the downregulation of glycolysis and upregulation of the TCA cycle in activated Tregs (Figure S5A and S5B). In terms of OXPHOS enrichment in the pathway analysis, Complexes I and IV were upregulated in freshly isolated Tregs, whereas Complex V was upregulated in activated Tregs (Figures S6A–S6C). These results are consistent with those of previous studies, which indicate that Tregs have reduced glycolysis and increases OXPHOS. As expected, RNA-seq revealed that activated Tregs had a distinct gene expression profile. However, activated Teffs and Tregs had similar metabolomic profiles, especially in terms of hydrophilic molecules, including carbohydrates and amino acids.

**Figure 1.**
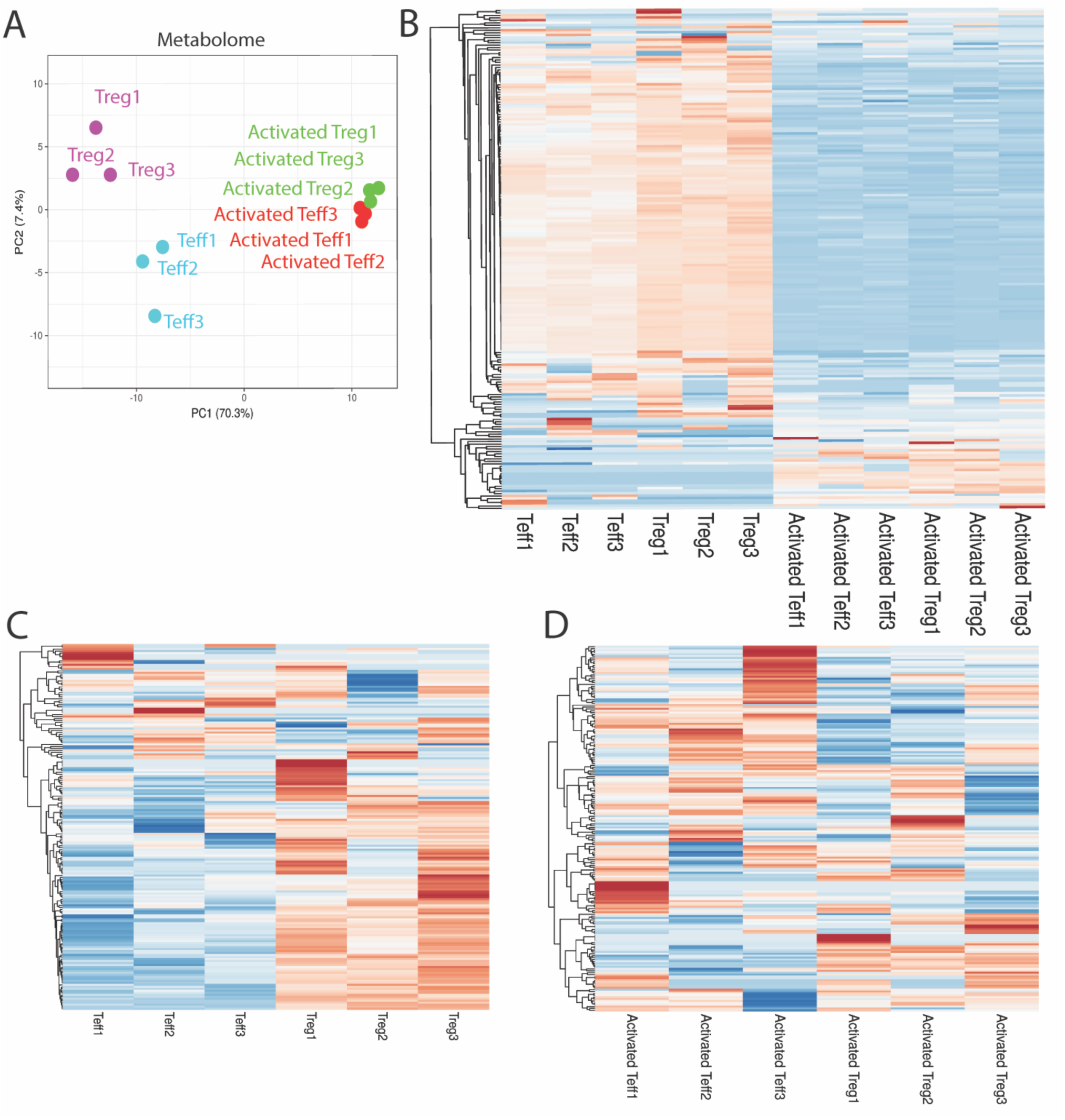
Freshly isolated Tregs have a distinctive metabolomic profile. (A–B) Metabolomic profile of freshly isolated and activated Tregs (n=3) shown as a PCA plot (A) and heatmap (B). (C–D) Metabolomic profile of freshly isolated Tregs (C) and activated Tregs (D) shown as a heatmap.

### Tregs had a unique lipidomic profile

The lipid profiles of Tregs were analyzed using lipidomic profiling. The lipidomics results are presented in Table S8. In the relative quantification, 611 lipids (23 lipid classes) were detected. Differentially accumulated lipids (DALs) in freshly isolated and activated Tregs are listed in Tables S9 and S10, respectively. Interestingly, freshly isolated Tregs exhibited a lipidomic profile including 8 DALs similar to that of freshly isolated Teffs (Figure 2A). Furthermore, *in vitro* TCR stimulation drastically altered the lipidomic profiles of the activated Tregs and Teffs including 75 DALs (Figure 2B). Although activated Teffs and Tregs had similar lipidomic profiles, unexpectedly, Tregs had increased triglycerides (TGs) containing unsaturated long-chain fatty acids but not medium-chain fatty acids (Figures 2C and 2D). Significant increases in unsaturated long-chain fatty acids were observed in TGs, such as TG(18:0_20:5_22:6), TG(20:4_20:5_22:6), TG(18:1_22:3_22:5), and TG(20:4_22:5_22:6) (*p*<0.01; Figure 2D). The overall TG distribution was not significantly altered in activated Tregs (Figures 3A and 3B). However, triacylglycerol lipase, which catalyzes TGs, was upregulated in activated Tregs but not in freshly isolated Tregs (Figures 3C and 3D). In contrast, unsaturated long-chain fatty acids were not upregulated in diglycerides (DGs) and free fatty acids, except for DG(18:1_20:1) and DG(18:0_22:6), but were significantly upregulated in activated Tregs (*p*<0.01) (Figures S3E and S3F).

**Figure 2.**
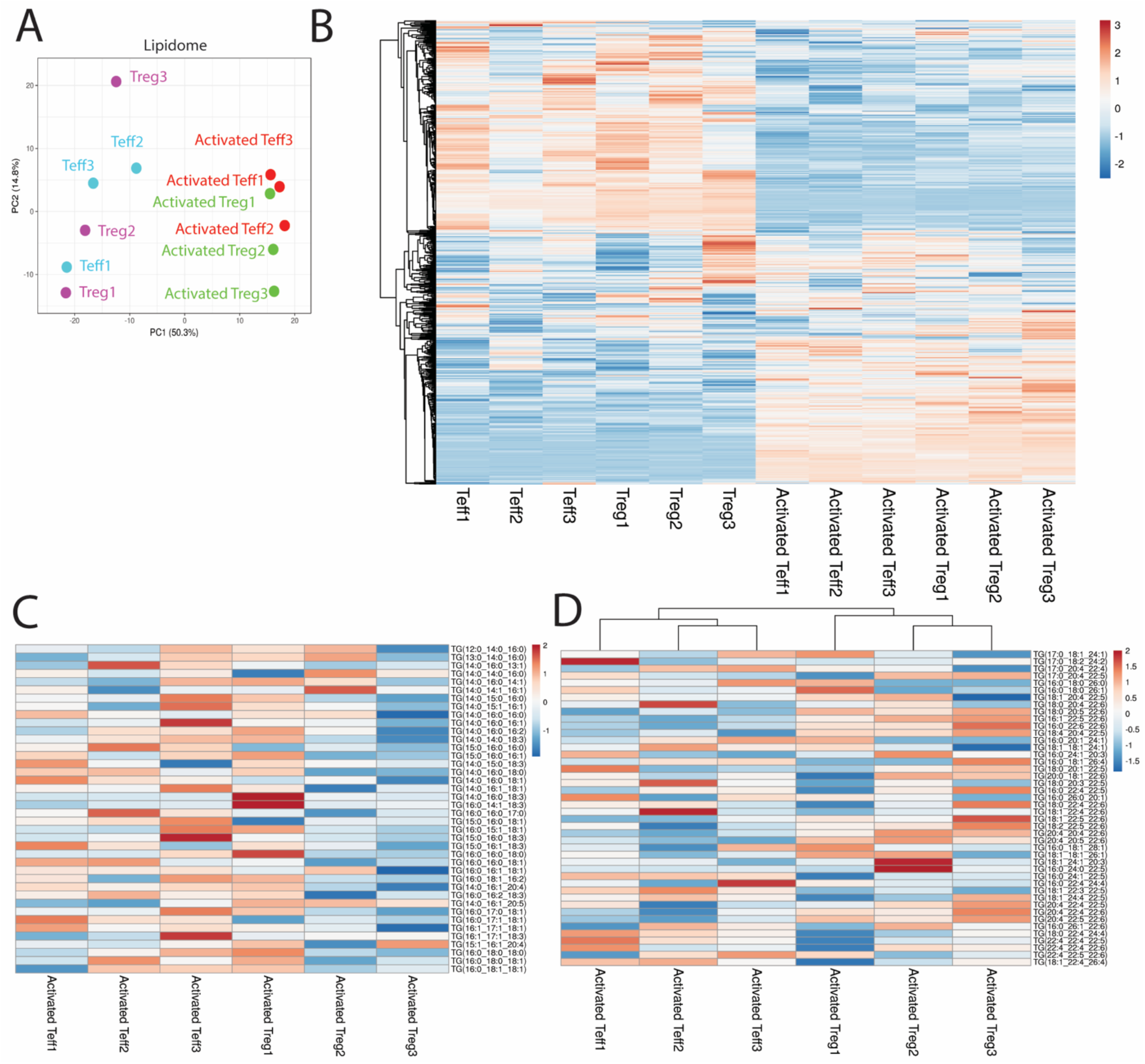
*In vitro* activated Tregs have a unique lipidomic profile. (A–B) Lipidomic profile of freshly isolated and activated Tregs (n=3) are shown as a PCA plot (A) and heatmap (B). (C–D) Lipidomic profile of activated Tregs derived TGs containing short-medium-chain (C) and long-chain fatty acids (D).

**Figure 3.**
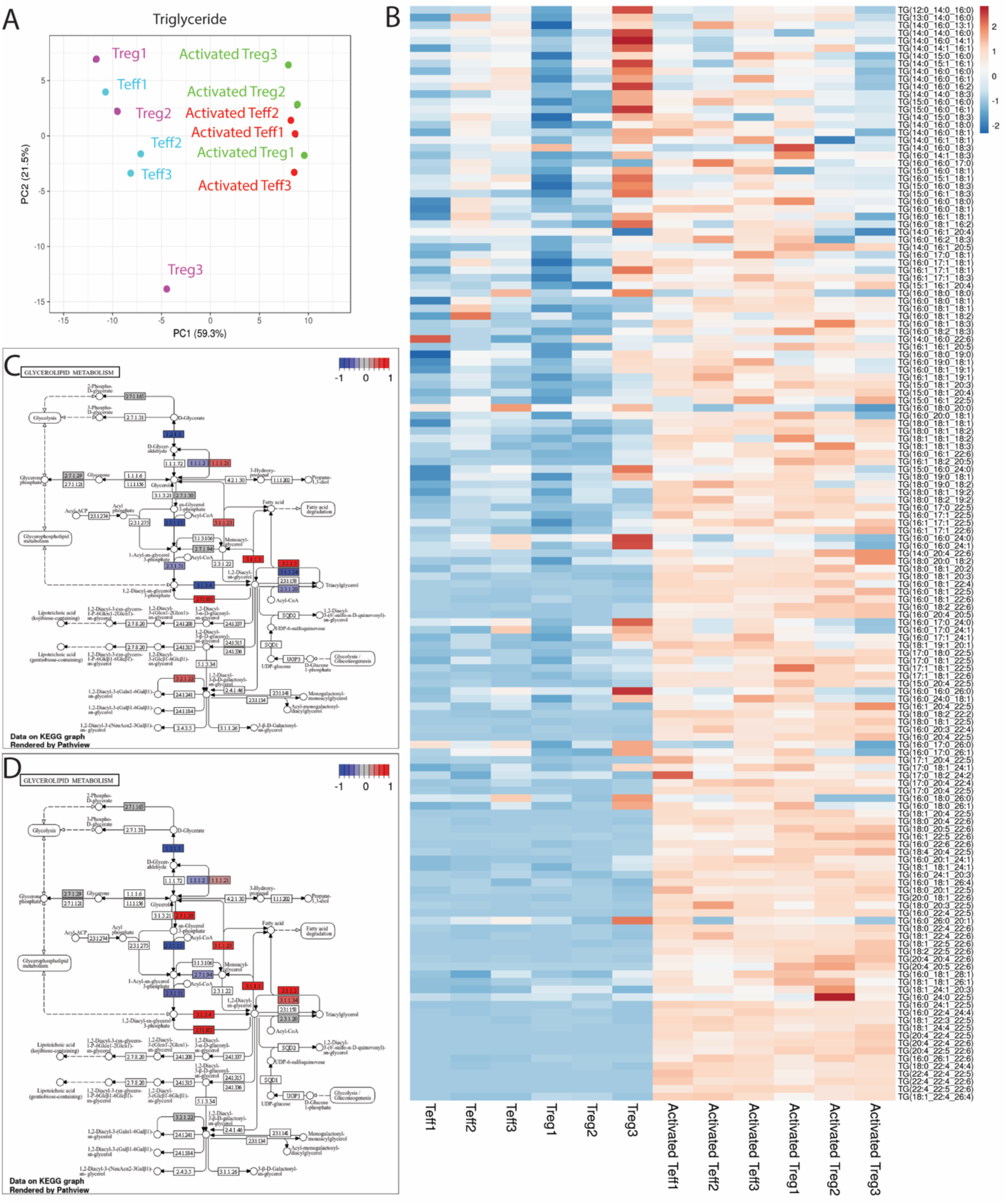
PUFA-rich TGs were upregulated in activated Tregs. (A–B) TG profile of freshly isolated and activated Tregs (n=3) shown as a PCA plot (A) and heatmap (B). (C–D) Gene expression profile of freshly isolated (C) and activated (D) Tregs a in the glycerolipid metabolism pathway.

TG(18:0_20:5_22:6) was also significantly upregulated in TGs; therefore, docosahexaenoic acid (DHA) was enriched in both DGs and TGs, but only in activated Tregs. In addition, RNA-seq identified a significant increase in the level of FABP4-5, which binds to fatty acids and regulates their transport, in activated Tregs (Figures 3G and 3H). To further evaluate TGs, the total counts of each fatty acid in DGs and TGs were determined. A significant increase in arachidonic acid (AA; C20:4), eicosapentaenoic acid (EPA; C20:5), and DHA (C22:6), but not in docosapentaenoic acid (DPA; C22:5) (*p*<0.05), was determined (Figure S7A). In contrast, AA and DPA did not increase, while DHA remained significantly increased in the DGs (*p*<0.05) (Figure S7B). EPA was not detected in DGs. Unsaturated long-chain fatty acids were not significantly altered in free fatty acids (Figure S7C). DHA enrichment was not detected in other lipid classes, namely phosphatidylcholine (PC), phosphatidylethanolamine (PE), phosphatidylglycerol (PG), phosphatidylinositol (PI), phosphatidylserine (PS), lysophosphatidylcholine (LPC), lysophosphatidylethanolamine (LPE), and cholesteryl esters (ChE). Collectively, these results reveal that compared with activated Teffs, activated Tregs contained omega-3 polyunsaturated fatty acid (PUFA)-DGs/TGs, but not other lipids, after TCR stimulation.

### Activated Tregs exhibited increased ceramide production

In addition to TGs, activated Tregs showed a significant increase in the accumulation of ceramides, including Cer(d18:0_16:0), Cer(d18:1_24:0), and Cer(d18:1_24:2) (*p*<0.01) (Figures 4A and 4B). While it was not possible to fully distinguish activated Tregs via metabolomic and lipidomic profiles, ceramide content could clearly distinguish activated Tregs from activated Teffs. RNA-seq also supported an increase in ceramide production in activated Tregs (Figures 4C and 4D). Moreover, the gene expression of ceramide-producing enzymes, including ceramide synthases (CERS1-6), increased in activated Tregs but not in freshly isolated Tregs. In addition to *de novo* synthesis, sphingomyelin synthase, which produces sphingomyelin and DGs, was upregulated in the activated Tregs. This may be associated with increased ceramide production and Treg-specific DG/TG signatures. Our data also confirms the increased ceramide content and gene expression profiles in activated Tregs.

**Figure 4.**
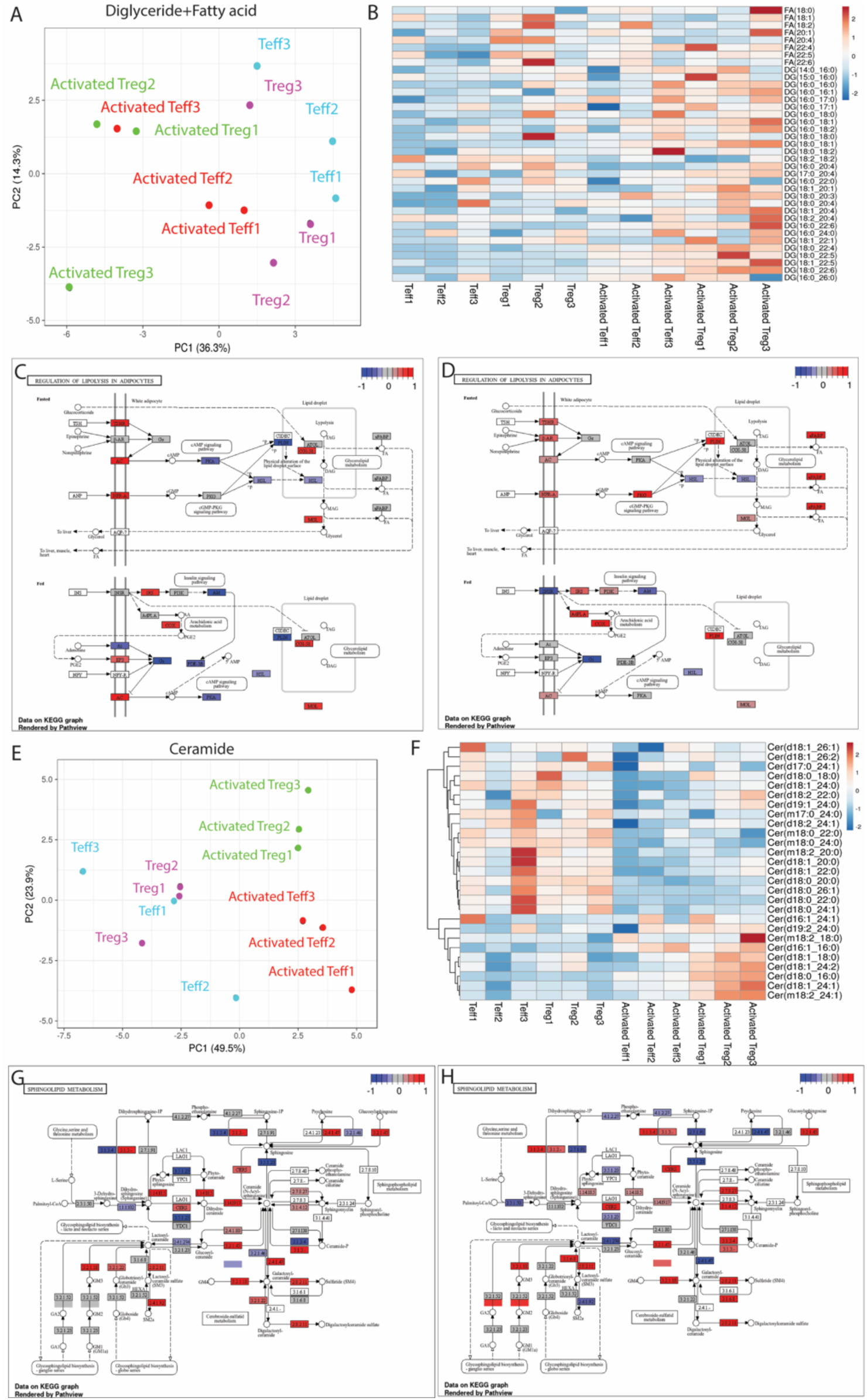
*In vitro* activated Tregs exhibit increased ceramide production. (A–B) Lipid profile of freshly isolated and activated Tregs (n=3) shown as a PCA plot (A) and heatmap (B). (C–D) Gene expression of freshly isolated (C) and activated (D) Tregs and in the lipolytic pathway. Ceramide accumulation shown as a PCA plot (D) and heatmap (E). Gene expression of freshly isolated (F) and activated (G) Tregs in the ceramide production pathway.

### Integrated transcriptomic, metabolomic, and lipidomic analyses reveal a unique lipid profile of Tregs upon activation

According to the results of the metabolomic and lipidomic profiling, freshly isolated Tregs had a unique metabolomic profile, whereas activated Tregs had a unique lipidomic profile upon TCR stimulation, characterized by DHA-rich DGs/TGs and enhanced ceramide production. As the metabolomic and lipidomic profiles were independently measured via mass spectrometry, the datasets were combined to assess the integrity of the hydrophilic and hydrophobic molecules. Combining metabolomic and lipidomic profiles as well as determining the metabolic status of the Tregs before and after TCR stimulation were possible (Figures 5A and 5B). Our results were consistent with the unique gene expression profiles of each subset before and after TCR stimulation. In contrast to metabolomic and lipidomic datasets, transcriptomic datasets contained >40000 transcripts and potentially interfered with the metabolic status. Therefore, direct combination of these datasets was not possible. To further investigate the relationship between transcripts and metabolites, transcriptomic, metabolomic, and lipidomic analyses were integrated and computationally analyzed simultaneously (Figure 5C). Interestingly, associations were found between transcripts-metabolites, such as TRIB2-TG(20:4_22:5_22:6), LILRA6-Cer(d18:0_16:0), HLA-DRB1-TG(17:0_20:4_22:5), TLR8-TG(16:0_22:6_22:6), MKKS-TG(16:0_20:4_22:5), and TMEM176B-TG(16:0_20:4_20:5). Among these, TRIB2, LILRA6, HLA-DRB1, TLR8, and TMEM176B were highly expressed in activated Tregs. In contrast, MKKS was preferentially expressed in Teffs, which is consistent with the results of previous studies (Preston et al., 2021).

**Figure 5.**
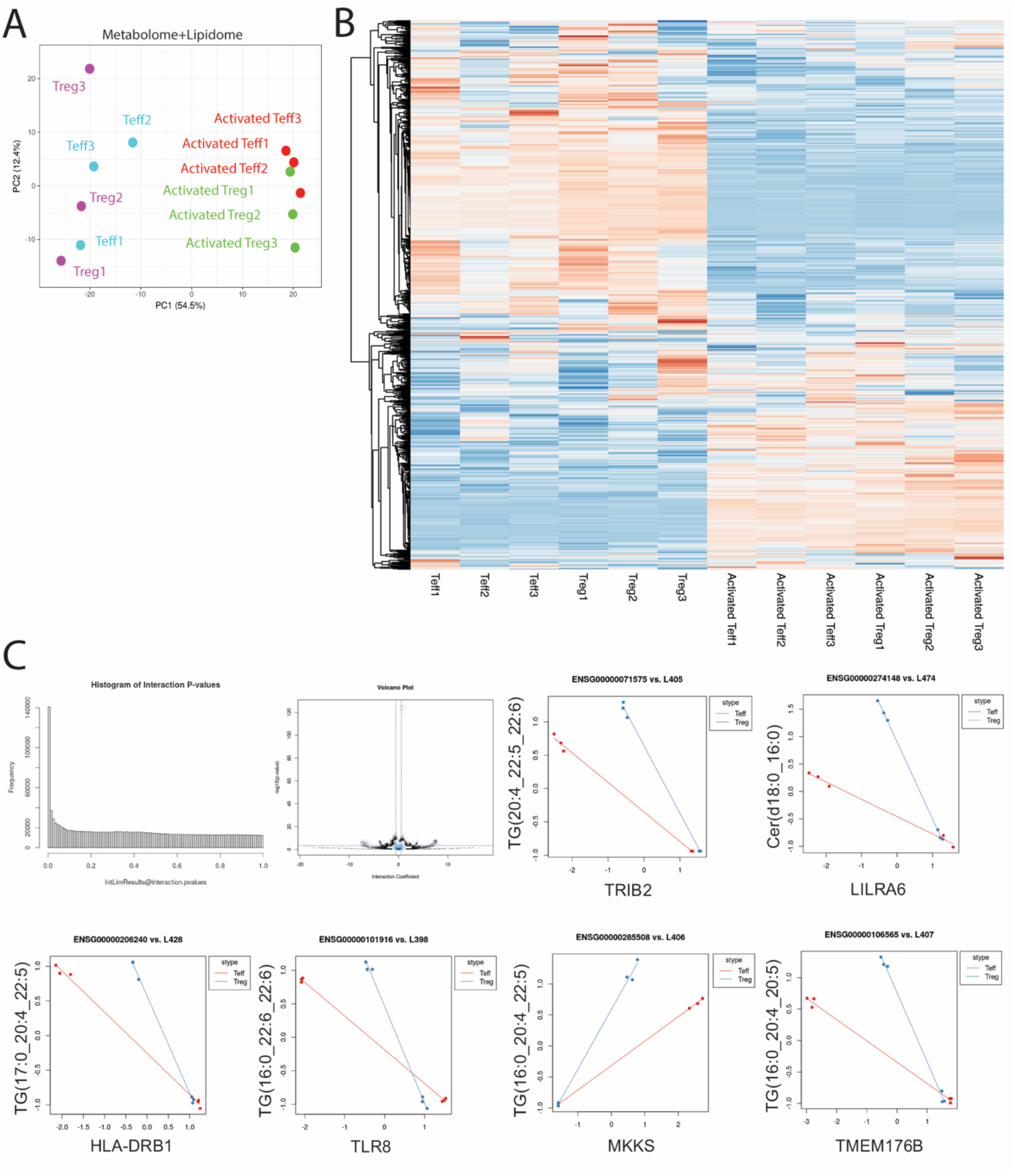
Integrated transcriptomic, metabolomic, and lipidomic analyses revealed a unique lipid profile of Tregs upon activation. (A–B) Combined metabolomic and lipidomic profiles shown as a PCA plot (A) and heatmap (B). (C) Integrated transcriptomic, metabolomic, and lipidomic analyses were performed using IntLIM (x-axis, transcript; y-axis, metabolites).

### Tregs had enriched lysosomes and mitochondria

To clarify the lysosomal changes in Tregs, the gene expression profiles of lysosomal membrane proteins (LAMP1 and LAMP2) and master transcription factors (TFE3 and TFEB) were analyzed. Upon activation, Tregs showed an increased expression of lysosomal proteins (Figures 6A and 6B). Moreover, LAMP1 and LAMP2 expression along with mitochondrial membrane potential (MT-1) were measured using flow cytometry. Gating strategies are shown in Figures S8A and S8B. LAMP-1 and LAMP-2 were significantly upregulated in activated Tregs compared with that in Teffs (Figure 6C), whereas mitochondrial membrane potential was not significantly elevated in freshly isolated Tregs (Figure 6D). To investigate the role of lysosomes and mitochondria in Tregs, we assessed mitochondrial and lysosomal morphology using immunohistochemistry (Figure 6E). While freshly isolated Teffs were not enriched in the lysosomes and mitochondria, activated Teffs were enriched in mitochondria upon TCR stimulation. Freshly isolated and activated Tregs were enriched in the lysosomes and mitochondria.

**Figure 6.**
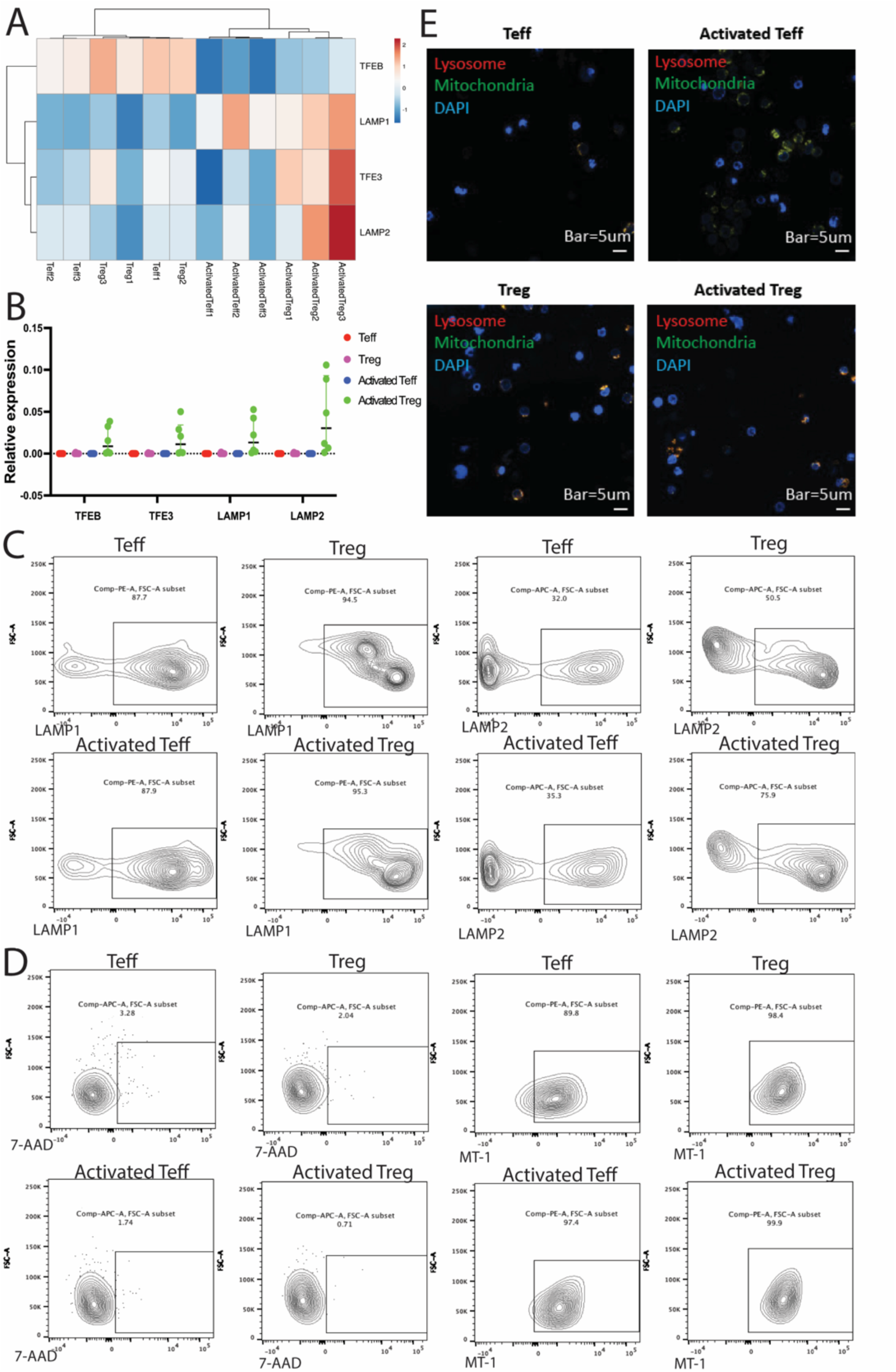
Tregs exhibited LAMP1/LAMP2 upregulation and enriched lysosomes. Gene expression related to lysosomes is shown by as RNA-seq (n=3) (A) and qPCR (n=6) (B). (C–D) Flow cytometry of activated Tregs (n=4) stained with LAMP1/2 (C) and mitochondrial membrane potential (D). (E) Live cell imaging for lysosomes and mitochondria.

### Tregs had upregulated PPAR-alpha/gamma and enriched peroxisomes

To elucidate the biological effect of increased levels of unsaturated long-chain fatty acids, we investigated the expression profiles of peroxisome-related genes. PPAR-alpha (PPARA) and PPAR-gamma (PPARG) were upregulated in the activated Tregs (Figure 7A). As PUFAs are taken up by fatty acid receptors (FFAR1 and FFAR4), FFAR expression was assessed using qPCR (Figure 7B). Freshly isolated Tregs expressed FFAR1/FFAR2/FFAR4, which take up PUFA upon activation. Activated Tregs also expressed FFAR1/FFAR2/FFAR3 upon TCR stimulation; however, their expression levels were similar to those in activated Teffs, despite the increased expression of PPARA/PPARG. Furthermore, we confirmed the upregulation of PPARG in both Teffs and Tregs upon activation (Figure 7C). In contrast, CD36 expression was not significantly altered in either Teffs or Tregs upon activation (Figure 7C). To clarify the role of peroxisomes in Tregs, peroxisomes were measured using immunohistochemistry. In the resting state, the Tregs had a similar peroxisomal morphology; however, activated Tregs exhibited peroxisome enrichment upon activation (Figure 7D). Collectively, Tregs were upregulated PPARA/PPARG upon activation and exhibited peroxisome enrichment. This molecular and organelle remodeling in activated Tregs might explain the metabolic alteration confirmed via integrated transcriptomic, metabolomic, and lipidomic analyses.

**Figure 7.**
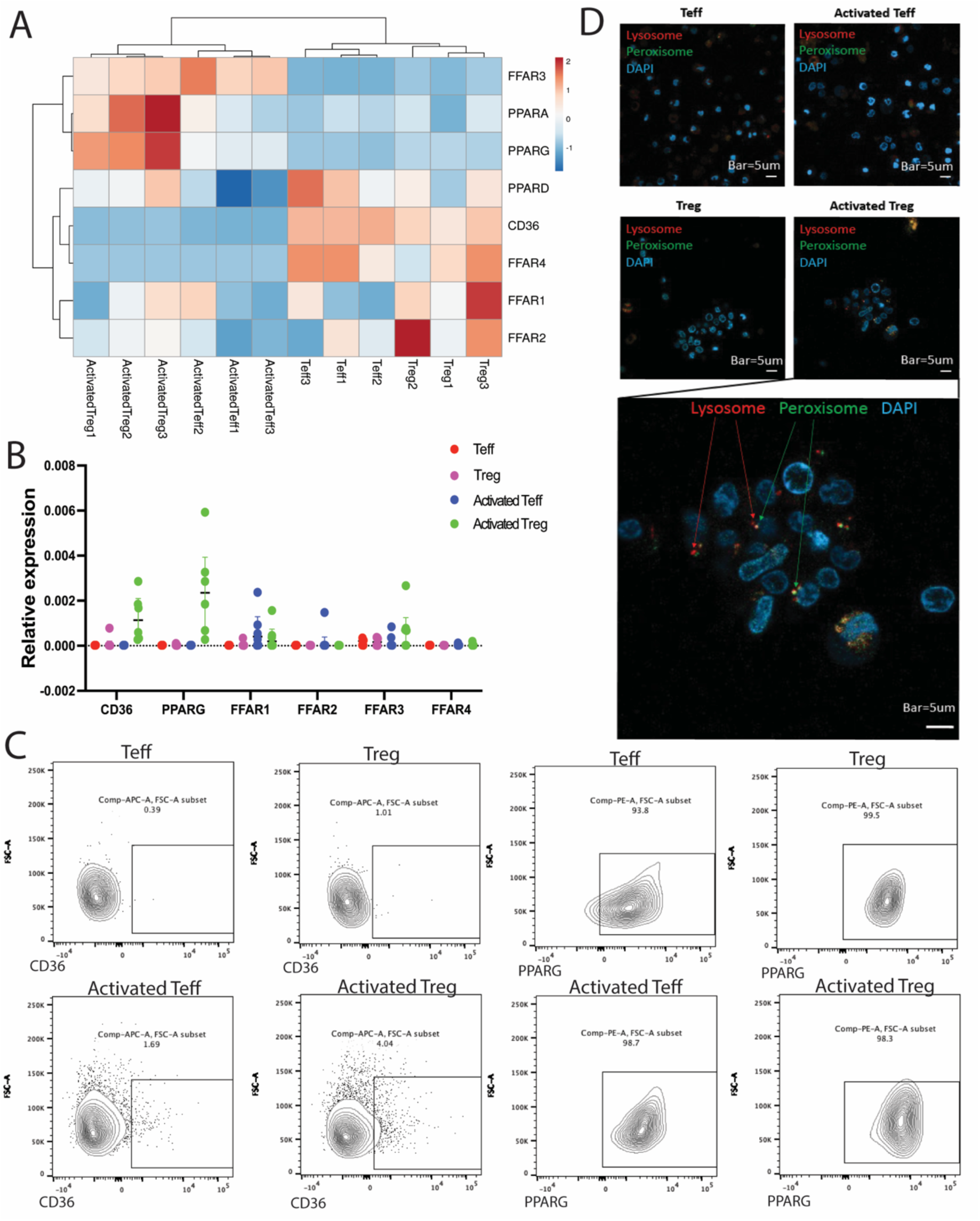
Tregs upregulated PPAR-gamma and had enriched peroxisomes. Gene expression related to fatty acid metabolism is shown by (A) RNA-seq (n=3) and (B) qPCR (n=6). (C) Flow cytometry of activated Tregs with regard to CD36/PPAR-gamma (n=4). (D) Live cell imaging for peroxisomes and lysosomes in freshly isolated and activated Tregs.

## Discussion

Our data indicate that activated Tregs had unique lipidomic profile, which included increased DGs/TGs consisting of long-chain unsaturated fatty acids, such as DHA. To support this, PPARA and PPARG were upregulated in Tregs.

Tregs play a central role in immune tolerance and homeostasis (Sakaguchi et al., 2010). Tregs have unique metabolomic properties, such as reduced glycolysis and increased oxidative phosphorylation (OXPHOS) compared with Teffs (Matias et al., 2021). Although transcriptomic and proteomic analyses of Tregs have been previously conducted (Bhairavabhotla et al., 2016; Procaccini et al., 2016), their metabolomic and lipidomic profiles have not yet been fully elucidated. Moreover, the studies focused on either transcriptomics or metabolomics, and only few combined different omics analyses. Metabolic alterations have been suggested to endow immune cells with robustness and functional properties, especially in the tumor microenvironment (Lim et al., 2021). These results support the need for further investigations of Tregs, including the process of lipid metabolism.

As Tregs mainly utilize OXPHOS for energy production, Tregs have also been shown to have enriched mitochondria (Beier et al., 2015). However, whether other endosomes, such as lysosomes and peroxisomes, are involved in this unique metabolic status remain unclear. Notably, Tregs exhibited an increased number of lysosomes and peroxisomes upon TCR stimulation. Importantly, metabolomic and lipidomic analyses indicated a similar metabolic status between Tregs and Teffs; however, gene expression analysis revealed a mild increase in the expression of genes related to lysosomes and peroxisomes. This might indicate that Tregs are prepared for activation even at baseline.

Unexpectedly, we found increased DHA in the DG/TG fractions, but not in the other lipid fractions. As synthesis of DHA in human cells is not possible, especially within short time periods (<24 h), DHA is most likely taken up by Tregs and combined with DGs or TGs. The functional role of DHA-containing DGs and TGs is not yet known; however, DHA-containing TGs are already available internationally as supplements. Convincing evidence for DHA-TGs remains lacking; however, Tregs, especially activated Tregs, may be enhanced by daily supplementation (Kim et al., 2018). In contrast, DHA deficiency or depletion is strongly associated with slow recovery from trauma and cirrhosis (Desai et al., 2014; Enguita et al., 2019). Treg dysfunction may not be the direct cause of these diseases, but could potentially affect or modulate their onset and progression via immune dysregulation. Moreover, fish are the major dietary sources of EPA and DHA, indicating the influence of dietary habits. Our data highlight the importance of healthy diet enriched in fish oils as a dietary source of omega-3 PUFA. Future clinical studies analyzing the association between PUFA intake and immunity may help clarify the beneficial effects of EPA and DHA intake or supplementation.

With regard to ceramide biosynthesis, Foxp3 has been suggested to inhibit Sgms1 in mice and this suppressive function could be maintained by increased ceramide via modulation of metabolic pathways (Apostolidis et al., 2016). In support of this, Sgms1 inhibition enhances Treg suppressive functions (Hollmann et al., 2016). This indicates that Tregs may have increased ceramide production, which may be relevant to their phenotype and function. Sphingomyelin synthase also produces DGs. Unlike DGs/TGs, which have a similar fatty acid profile, including enriched DHA, ceramides do not contain PUFAs. Therefore, PUFAs such as EPA and DHA are most likely taken up by the cells upon activation. How ceramide production and accumulation modulate Treg function warrants further study.

Recent research highlights the interplay between omega-3 PUFA, Tregs, and ceramide pathways in modulating immune responses. In mice, omega-3 PUFA can reverse the inflammatory effects of a Western diet by reducing ceramide levels and enhancing Treg populations, thereby improving immune function (Camacho-Munoz et al., 2022). Similarly, dietary components, including omega-3 PUFA, are important in maintaining Treg homeostasis and preventing chronic inflammation (Issazadeh-Navikas et al., 2012). In the mice, EPA induces Tregs through a PPARγ-dependent mechanism, suggesting potential therapeutic applications for autoimmune diseases (Iwami et al., 2011). Lastly, the broader molecular mechanisms by which omega-3 fatty acids exert beneficial health effects, including their regulatory roles in immune and metabolic pathways, were shown previously (Seo et al., 2005). Collectively, these studies underscore the significance of omega-3 fatty acids in enhancing Treg function and mitigating inflammation. The unique organelle composition of immune cells is associated with their functions. Mitochondrial function is highly associated with immune function via energy production (Beier et al., 2015; Field et al., 2020). Moreover, lysosomes are related to immune functions, including Treg suppression, and potentially interact with the mitochondria (Xia et al., 2022). Compared with mitochondria and lysosomes, peroxisomes have rarely been described in the immune system; however, they have been recognized as immune regulators, especially through lipid metabolism (Di Cara et al., 2023). Our study identified the transcriptomic-metabolism-organelle axis to understand cell biology.

Overall, our results showed the importance of lipid metabolism, including omega-3 long-chain PUFA-DGs/TGs and ceramide biosynthesis, in Treg function. Compared with the transcriptome and metabolome, the lipidomic profile of the immune system is now well studied. However, lipid metabolism may provide the metabolic regulation of immunity. In addition, integrated transcriptomic, metabolomic, and lipidomic profiles may help identify novel transcriptomic/metabolic profiles of immune cells.

## STAR Methods

### PBMC isolation

Frozen PBMCs (Precision of Medicine; FUJIFILM Wako Pure Chemical, Japan) were stored in liquid nitrogen. Upon thawing, Roswell Park Memorial Institute medium (FUJIFILM Wako Pure Chemical) supplemented with 10% fetal bovine serum (Gibco, MA, USA) and 1% penicillin-streptomycin (Gibco) was added to the cryovials, and the resuspended samples were transferred to a 15-mL tube. Next, 50 U/mL Benzonase (MilliporeSigma, MA, USA) was added to the frozen samples resuspended in culture medium. The thawed PBMCs were centrifuged at 400 × *g* for 5 min at room temperature. After carefully removing the supernatant, the cells were resuspended in culture medium for subsequent antibody staining.

### Cell sorting

PBMCs were incubated with Fc Block (BD Biosciences, NJ, USA) for 15 min on ice and then stained with an antibody cocktail consisting of anti-CD3, anti-CD4, anti-CD25, and anti-CD127 antibodies (BioLegend, CA, USA) for 30 min at 4 °C. After staining, the cells were washed thrice with FACS buffer (phosphate-buffered saline [PBS] supplemented with 0.5 mM EDTA and 1% bovine serum albumin). The cells were resuspended in FACS buffer at a concentration of 1e6 cells/mL and immediately processed for cell sorting. Tregs and Teffs were sorted in the culture medium via FACS on FACSARIA III (BD Biosciences), according to the gating strategy shown in Figure S1B. After sorting, the cells were centrifuged, and cell numbers and viability were determined using Countess III (Thermo Fisher Scientific, Waltham, MA, USA). Flow cytometric data were analyzed using FlowJo software (FlowJo, LLC, OR, USA). List of antibodies used in this study was provided in Table S11.

### In vitro culture

Sorted Tregs and Teffs (concentration of 1e6 cells/mL) were resuspended in a culture medium supplemented with 100 U/mL recombinant human IL-2 (PeproTech, NJ, USA) and 25 µL/mL Immunocult Human CD3/CD28/CD2 T Cell Activator (STEMCELL Technologies, BC, Canada). After 16-24 h of incubation at 37 °C, the cells were collected and RNA/metabolites were extracted.

### Flow cytometry

Teffs and Tregs were analyzed using flow cytometry to assess the metabolic status and the status of organelles. Similar to what was done for cell sorting, the cells were incubated with Fc Block for 15 min on ice, stained with antibody cocktails consisting of anti-CD3, CD4, CD25, CD36, CD127 antibodies, and subjected to 7-AAD live dead staining for 30 min at 4 °C. After staining, the cells were washed thrice with FACS buffer. The cells were fixed and permeabilized using an e-Bioscience FOXP3 Staining Buffer kit (Thermo Fisher Scientific). The cells were then stained with an antibody cocktail containing FOXP3, LAMP1, LAMP2, and PPARG. The mitochondrial membrane potential was measured using the MT-1 MitoMP Detection Kit (Dojindo Laboratories, Japan) according to the manufacturer’s protocol. The stained cells were analyzed via FACS on FACSARIA III. Flow cytometric data were analyzed using FlowJo software. List of antibodies used in this study was provided in Table S11.

### RNA isolation

Cells (1e6) were washed twice with PBS, and RNA was isolated using an RNeasy Micro Kit (QIAGEN, Germany). RNA quality and quantity were measured using an RNA 6000 Nano kit on a Bio Analyzer (Agilent Technologies, CA, USA). RNA samples with RIN>8.0 were used for RNA-seq.

### qPCR

The isolated mRNA was reverse transcribed to cDNA using ReverTra Ace qPCR RT Master Mix (Toyobo Co., Ltd., Osaka, Japan). Real-time PCR was conducted on QuantStudio 5 (Thermo Fisher Scientific) using a TaqMan gene expression assay (HPRT, TFEB, TFE3, LAMP1, LAMP2, CD36, PPARG, FFAR1, FFAR2, FFAR3, and FFAR4; Thermo Fisher Scientific). The expression of target genes is presented as relative expression normalized to the housekeeping gene (HPRT) expression. List of TaqMan probes used in this study was provided in Table S12.

### RNA-seq

RNA-seq was performed by Rhelixa Co., Ltd. (Tokyo, Japan). Briefly, mRNA was purified from total RNA samples using the Poly-A selection method [NEBNext® Poly(A) mRNA Magnetic Isolation Module; New England Biolabs, MA, USA], and single-stranded cDNA was synthesized via reverse transcription using this as a template (NEBNext® UltraTM ll Directional RNA Library Prep Kit; New England Biolabs). The single-stranded cDNA was used as a template for second-strand cDNA synthesis using nucleoside triphosphates containing dUTP instead of dTTP. Both ends of the resulting double-stranded DNA were blunted and phosphorylated, followed by the creation of 3’-dA overhangs and ligation of a sequencing adapter containing dUTP. The dUTP in the adapter and second-strand cDNA were selectively cleaved using the USER enzyme to obtain a template for a library synthesized from single-stranded DNA with directional information (strand-specific library). Base sequences of the library-prepared samples were obtained using a next-generation sequencer (Illumina NovaSeq 6000, 150bp×2 paired-end). In particular, 4 G bases per sample were sequenced and 26.7 M reads per sample (13.3 M pairs) were obtained. Raw sequence reads were quantified using Salmon (Patro et al., 2017) and further analyzed using iDep software (Ge et al., 2018). Raw sequence data have been deposited at ArrayExpress under accession number E-MTAB-14707.

### Metabolomic profiling

Metabolomic profiling was performed by the Human Metabolome Technologies Co., Ltd. (Yamagata, Japan). After culturing Tregs, the medium was removed, and the cells were washed twice with mannitol (FUJIFILM Wako Pure Chemical). Methanol (FUJIFILM Wako Pure Chemical) solution was added and stirred, followed by Milli-Q water containing 10 μM of an internal standard (HMT, patent no. 6173667) was added and stirred. The mixture was centrifuged (2,300 × *g*, 4 °C, 5 min) (patent no. 6173667), after which the extract was transferred to an ultrafiltration tube (Ultrafree MC PLHCC, HMT, centrifugal filter unit: 5 kDa). The extract was centrifuged (9,100 × *g*, 4 °C) and subjected to ultrafiltration. The filtrate was dried and dissolved in Milli-Q water for measurements. In this experiment, measurements were performed in cation mode and anion mode by Agilent CE-TOFMS system (Agilent Technologies). Peaks detected by CE-TOFMS were automatically extracted using the automatic integration software MasterHands ver.2.19.0.2 (developed by Keio University) with a signal-to-noise (S/N) ratio of 3 or more, and mass-to-charge ratio (m/z), peak area, and migration time (MT) were obtained. The peak area values obtained were converted to relative area values using the following formula.

Relative area value = (Area value of the target peak)/ (Area value of the internal standard substance × sample amount)

In addition, these data include adduct ions such as Na+ and K+, and fragment ions such as dehydrated and deammonium ions, so these molecular weight-related ions were deleted. However, since there are also substance-specific adducts and fragments, it was not possible to examine all of them. For the peaks examined, matching and alignment of peaks between each sample was performed based on the m/z and MT values. The detected peaks were matched and searched against all substances registered in the HMT metabolite library based on the m/z and MT values. The search tolerance was set to ± 0.5 min for MT and ± 10 ppm for m/z.

Mass error (ppm) = (Actual value – theoretical value)/ Actual value×10^6^

For substances with molecular weights of 100 or less and some substances expected to be present in large amounts in the sample, the tolerance was set wider than the above conditions and the search was performed. In addition, if the candidates could not be narrowed down and the same candidate metabolite was assigned to multiple peaks, a branch number was added and the list was displayed. The target metabolites were analyzed. The calibration curve was made using peak areas corrected by the internal standard substance, and the concentration was calculated for each substance with a single calibration of 100 μM (internal standard substance 200 μM). Metabolomic data have been deposited at Metabolomics Workbench (ST003626).

### Lipidomic profiling

Nontargeted lipidomic profiling was performed by Lipidome Lab. Co., Ltd. (Akita, Japan). Briefly, a cell suspension was prepared by adding methanol was added to frozen cell pellets. Total lipids were extracted from the entire suspension (equivalent to 1 × 10-6 cells) using a partially modified Bligh and Dyer method, which is a liquid-liquid distribution method that uses chloroform, methanol, and water. The extracted lipid fraction was dried with nitrogen gas, re-dissolved in 1 mL of methanol, and transferred to a measurement vial for analysis. Liquid chromatography (LC)-electrospray ionization-MS/MS analysis was performed by using Q-Exactive Plus mass spectrometer with an UltiMate 3000 LC system (Thermo Fisher Scientific). Samples were separated on L-column3 C18 metal-free column (2.0 µm, 2.0 mm × 100 mm i.d.) at 40°C using a gradient solvent system: mobile phase A (isopropanol/methanol/water (5/1/4 v/v/v) supplemented with 5 mM ammonium formate and 0.05% ammonium hydroxide (28% in water))/mobile phase B (isopropanol supplemented with 5 mM ammonium formate and 0.05% ammonium hydroxide (28% in water)) ratios of 60%/40% (0 min), 40%/60% (0-1 min), 20%/80% (1-9 min), 5%/95% (9-11 min), 5%/95% (11-22 min), 95%/5% (22-22.1 min), 95%/5% (22.1-25 min), 60%/40% (25-25.1 min) and 60%/40% (25.1-30 min). The injection volume was 10 µl and the flow rate was 0.1 mL/min. For the peaks examined, matching and alignment of peaks between each sample was performed based on the m/z and MT values. Lipid Search 5. 1 (Mitsui Knowledge Industry Co., Ltd., Tokyo, Japan), a lipid identification software, was used to identify lipid molecular species and align the measured samples. The correction value was calculated using the following equation (total area correction): Correlation value = (area of detected peak) / (total area of detected peak). A comprehensive lipidomic analysis of the samples was performed using ion chromatography/electrospray ionization/mass spectrometry. The obtained m/z values and fragment ions were compared with those in the database registered in the Lipid Search, and the number of molecular species that met the adoption criteria (our nondisclosed indicators) was 23 classes and 611 molecular species. The peak area of each identified molecule was corrected using the total area value (sum of the detected area values). Additionally, the relative values were used to correct the peak areas. Lipidomic data have been deposited at Metabolomics Workbench (ST003627).

### Integrated transcriptomic, metabolomic, and lipidomic analyses

Metabolomic and lipidomic data were combined and analyzed together because of their similarity in data size and format (relative quantification by mass spectrometry). To combine metabolomic and lipidomic data with transcriptomic data, integration was performed using IntLIM (Siddiqui et al., 2018). Significantly correlated transcriptional metabolites were identified after data integration.

### Live cell imaging

Freshly isolated Tregs/Teffs were resuspended in culture medium and incubated overnight with CellLight Lysosome-RFP, CellLight Peroxisome-GFP, and MitoTracker Green FM (Thermo Fisher Scientific). The following day, the cells were washed thrice with fresh medium and resuspended in PBS supplemented with Hoechst 33342 (Dojindo Laboratories). Cell images were acquired using an LSM980 with Airyscan2 (Zeiss, Germany) and analyzed using Zen software (Zeiss).

### Statistical analysis

GraphPad Prism Software (ver9.3.1; GraphPad Software, La Jolla, CA, USA) was used for statistical analyses. All statistical analyses of the metabolome and lipidome were performed using paired *t*-test. Principal component analysis and heatmap were performed and generated, respectively, using ClustVis (Metsalu and Vilo, 2015). Statistical significance was set at *P* < 0.05.

## Ethical approval

The study protocol was approved by the Institutional Review Board of the Jikei University School of Medicine (#36-304(12417)).

## Limitations of the study

Owing to the limited number of Tregs from each PBMC donor, multiple donors were pooled for metabolomic and lipidomic analyses. The number of samples was limited; however, the acquired values represented the average amount across different donors.

## Data availability

All data supporting our conclusions are mentioned in this article and in the Supporting Information.

## Supporting information

This article contains supporting information.

## Acknowledgments

We express our gratitude to Dr. Koji Hayashizaki (Department of Bacteriology, Jikei University School of Medicine) and Dr. Yuki Kinjo (Department of Bacteriology, Jikei University School of Medicine) for granting access to and providing technical support for BD Aria III.

## Authors’ contribution

Yohei Sato: Conceptualization; Yohei Sato and Misa Yura: Methodology; Yohei Sato: Formal analysis and investigation; Yohei Sato: Writing–original draft; Yohei Sato, Alana Chandler and Yamato Hanawa: Writing–review and editing; Akihito Tsubota: Supervision.

## Declaration of Generative AI and AI-assisted technologies in the writing process

Generative AI and AI-assisted technologies were not used in the writing process.

