## Supplemental Figures for "Integrated transcriptomic, metabolomic, and lipidomic analyses reveal a unique lipid profile of regulatory T cells upon activation"

**Supplementary Figures/Legends**

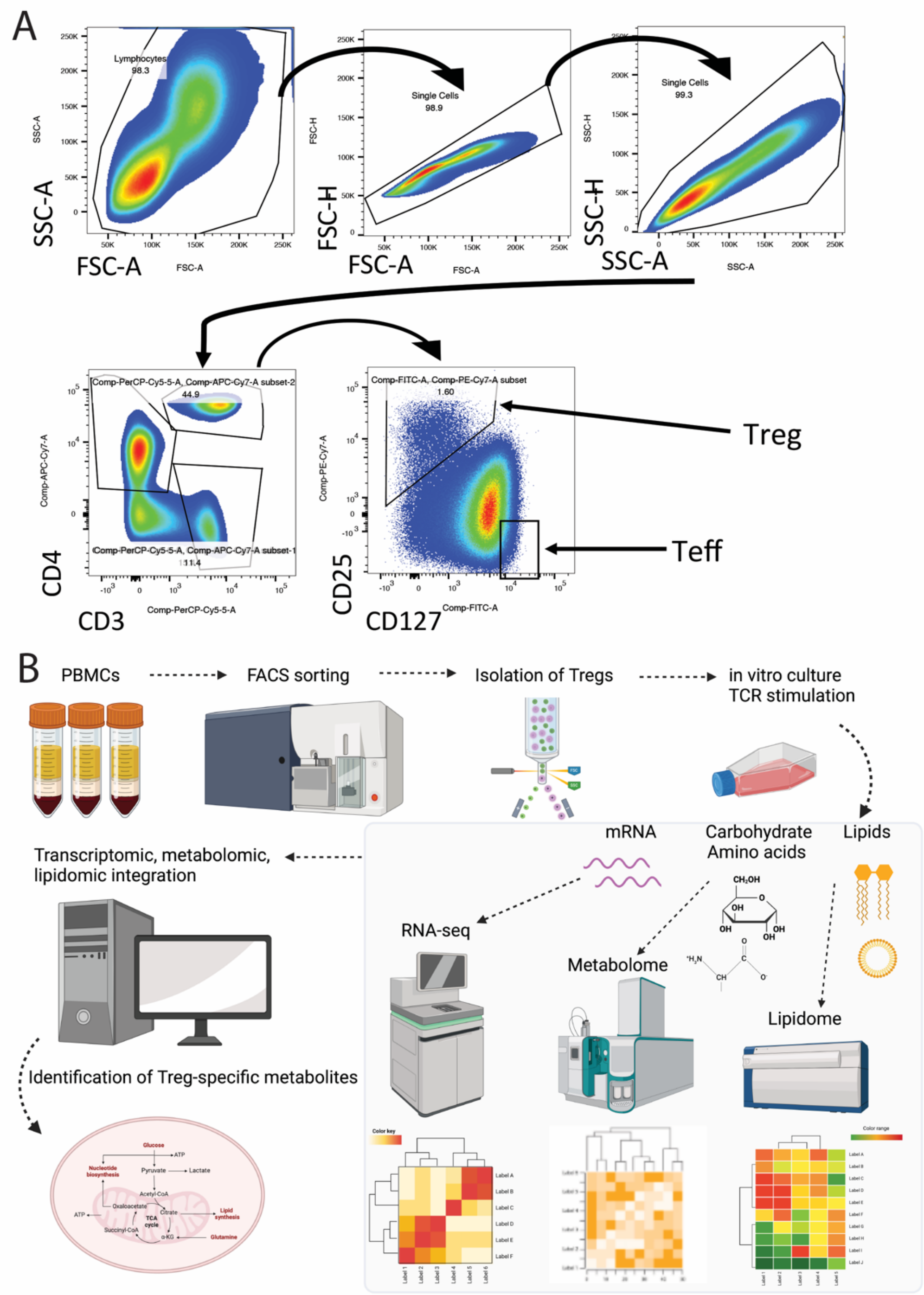


**Figure S1 Tregs can be isolated from PBMCs and pooled for transcriptomic, metabolomic, and lipidomic analyses.** (A) Gating strategy of Treg/Teff isolation. (B) Experimental and analytical design of the present study.

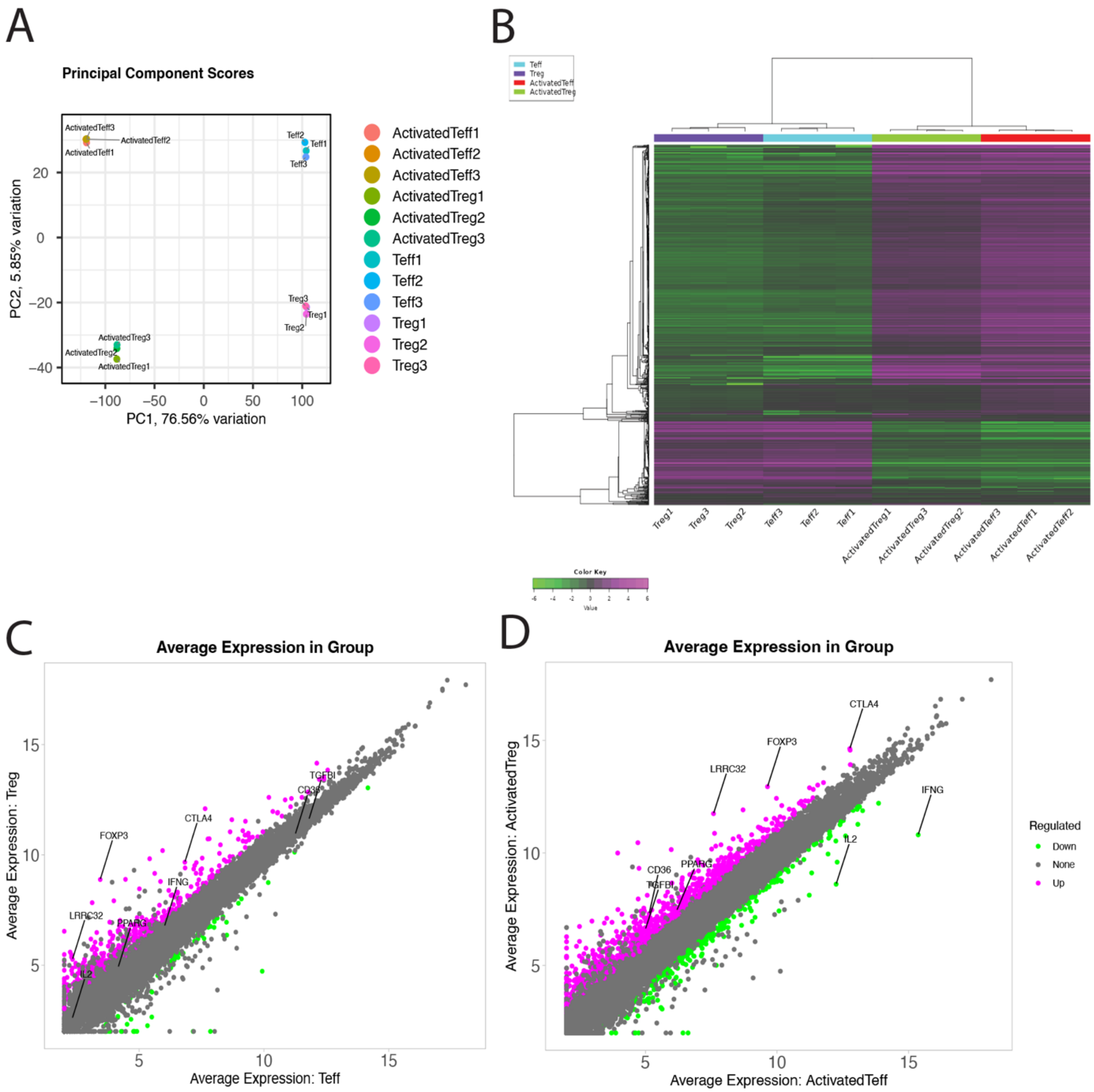


**Figure S2 Tregs have a distinct transcriptomic profile.** (A–B) Gene expression of freshly isolated and activated Teffs/Tregs shown as a PCA plot (A) and heatmap (B). (C–D) Gene expression of freshly isolated (C) and activated (D) Tregs shown as scatter plots.

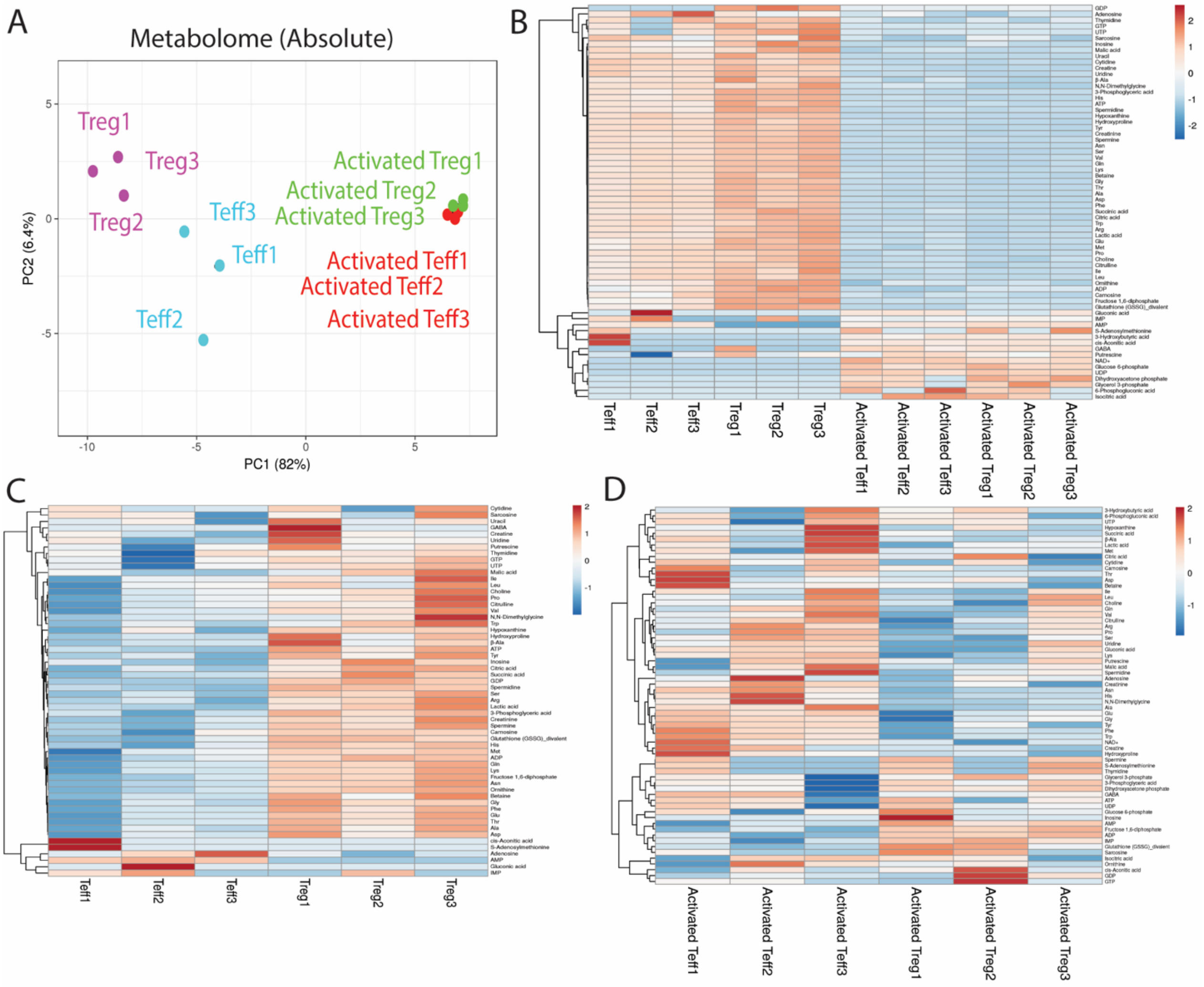


**Figure S3 Absolute measurement of the metabolomic profile.** (A–B) Absolute quantifications of metabolites shown as a PCA plot (A) and heatmap (B). (C–D) Abundance of each metabolite shown as a heat map for the freshly isolated (C) and activated (D) Tregs.

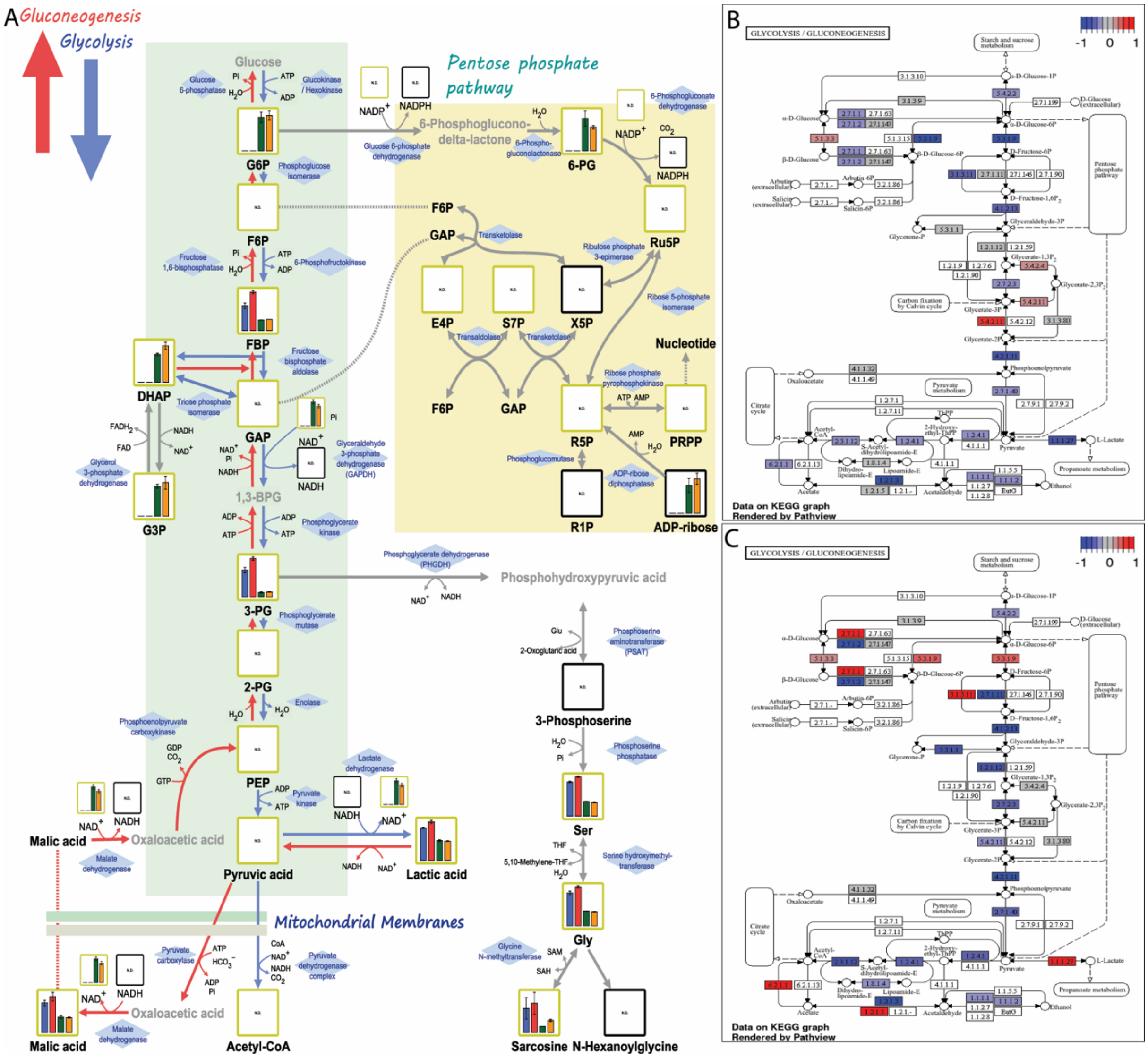

**Figure S4 Glycolytic pathway was not activated in Tregs.** (A) Each metabolite in the glycolytic pathway is shown in the metabolic map. (B) Expression of genes related to the glycolytic pathway was analyzed in freshly isolated (C) and activated (D) Tregs.

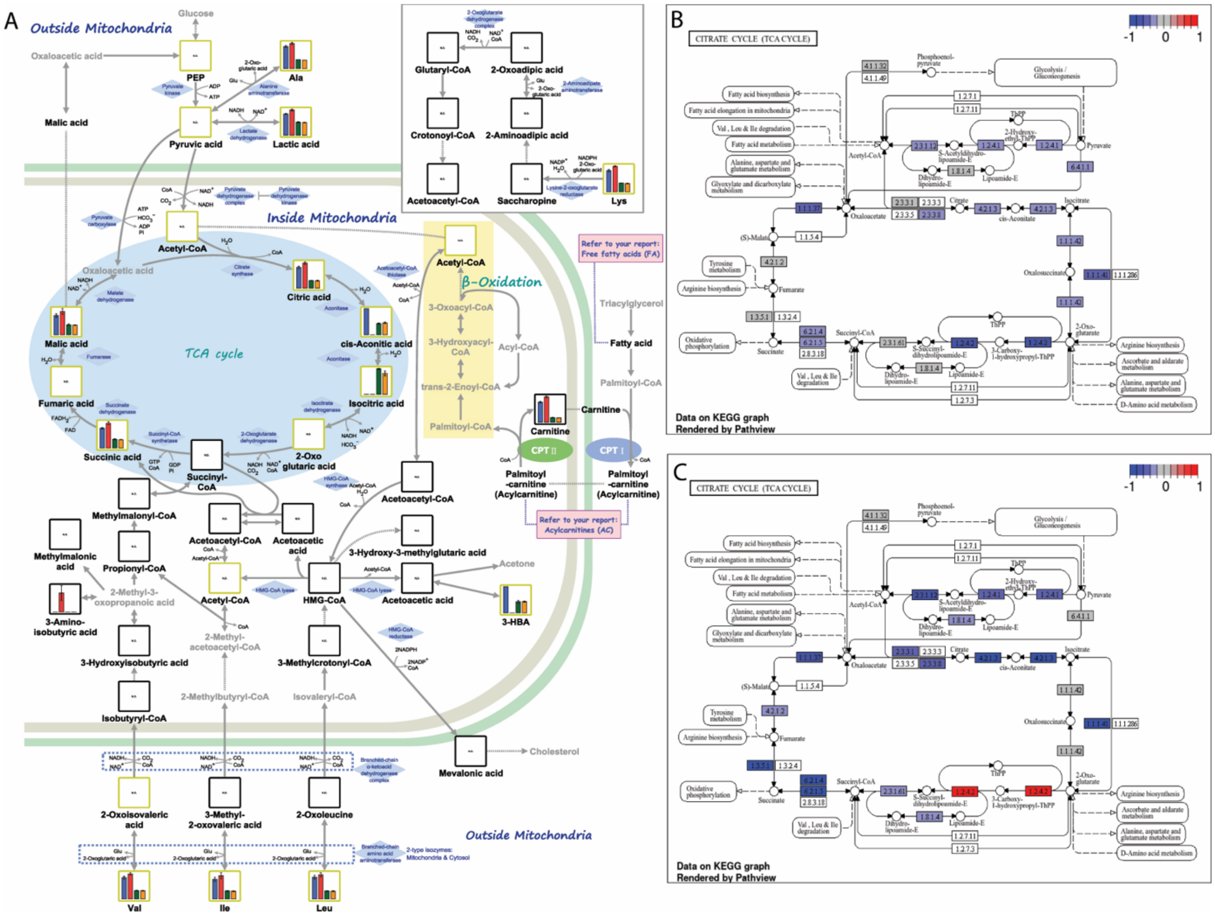


**Figure S5 TCA cycle was not activated in Tregs.** (A) Each metabolite in the TCA cycle is shown in the metabolic map. (B) Expression of genes related to the TCA cycle was analyzed in freshly isolated (C) and activated (D) Tregs.

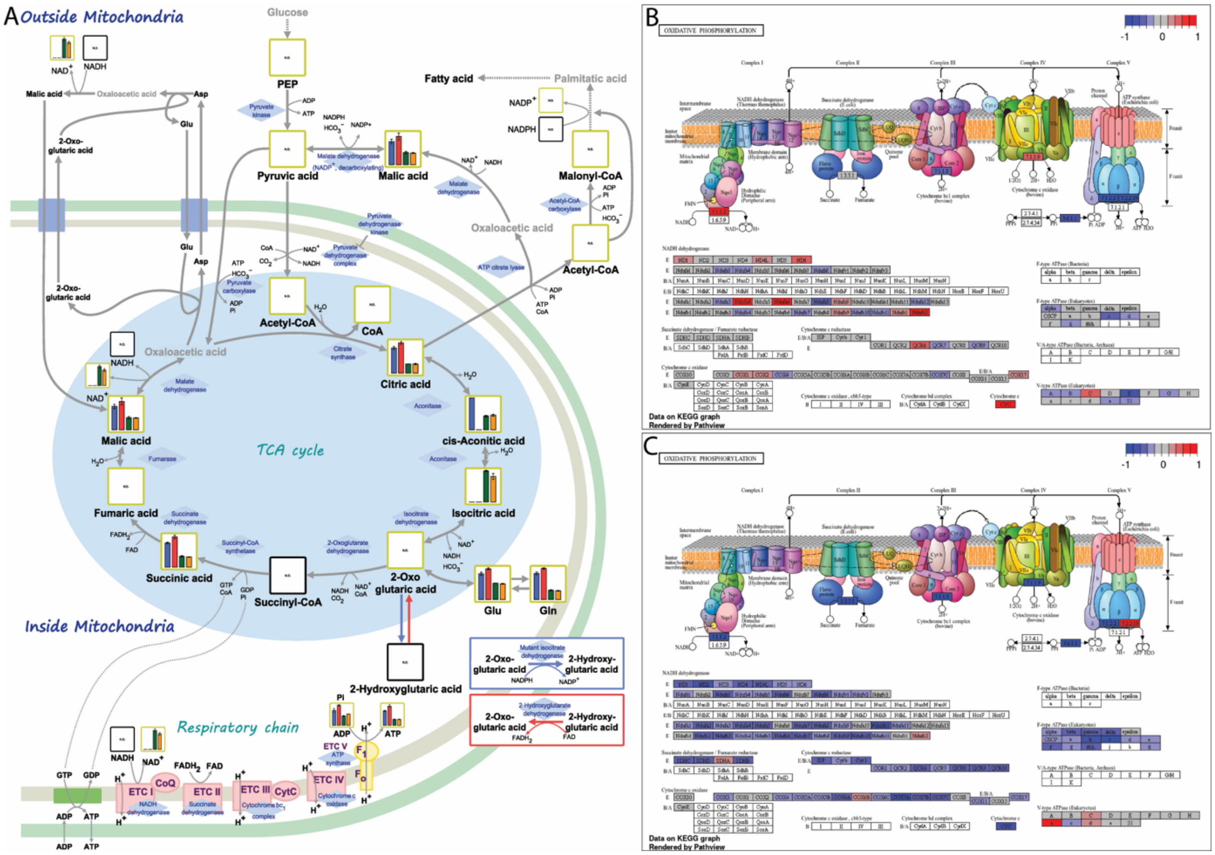


**Figure S6 OXPHOS pathway was partially activated in Tregs.** (A) Each metabolite in OXPHOS is shown in the metabolic map. (B) Expression of genes related to OXPHOS was analyzed in freshly isolated (C) and activated (D) Tregs.

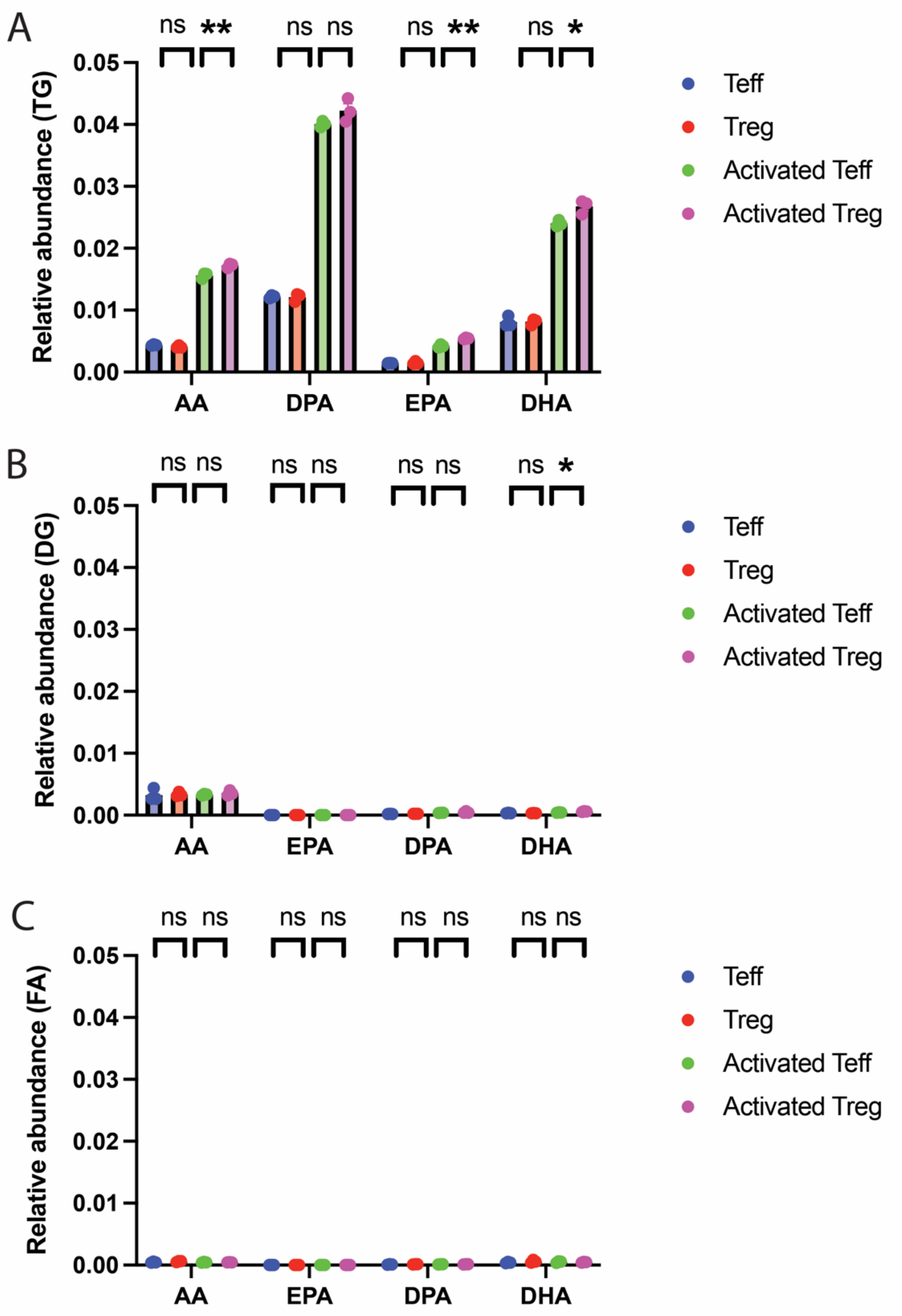


**Figure S7 Omega-3 PUFA were enriched in the DGs and TGs in Tregs upon activation.** (A–B) Long-chain fatty acids (AA, DPA, EPA, and DHA) were analyzed in TGs (A) and DGs (B) for each cell condition (n=3). (C) Long-chain free fatty acids for each cell condition (n=3).

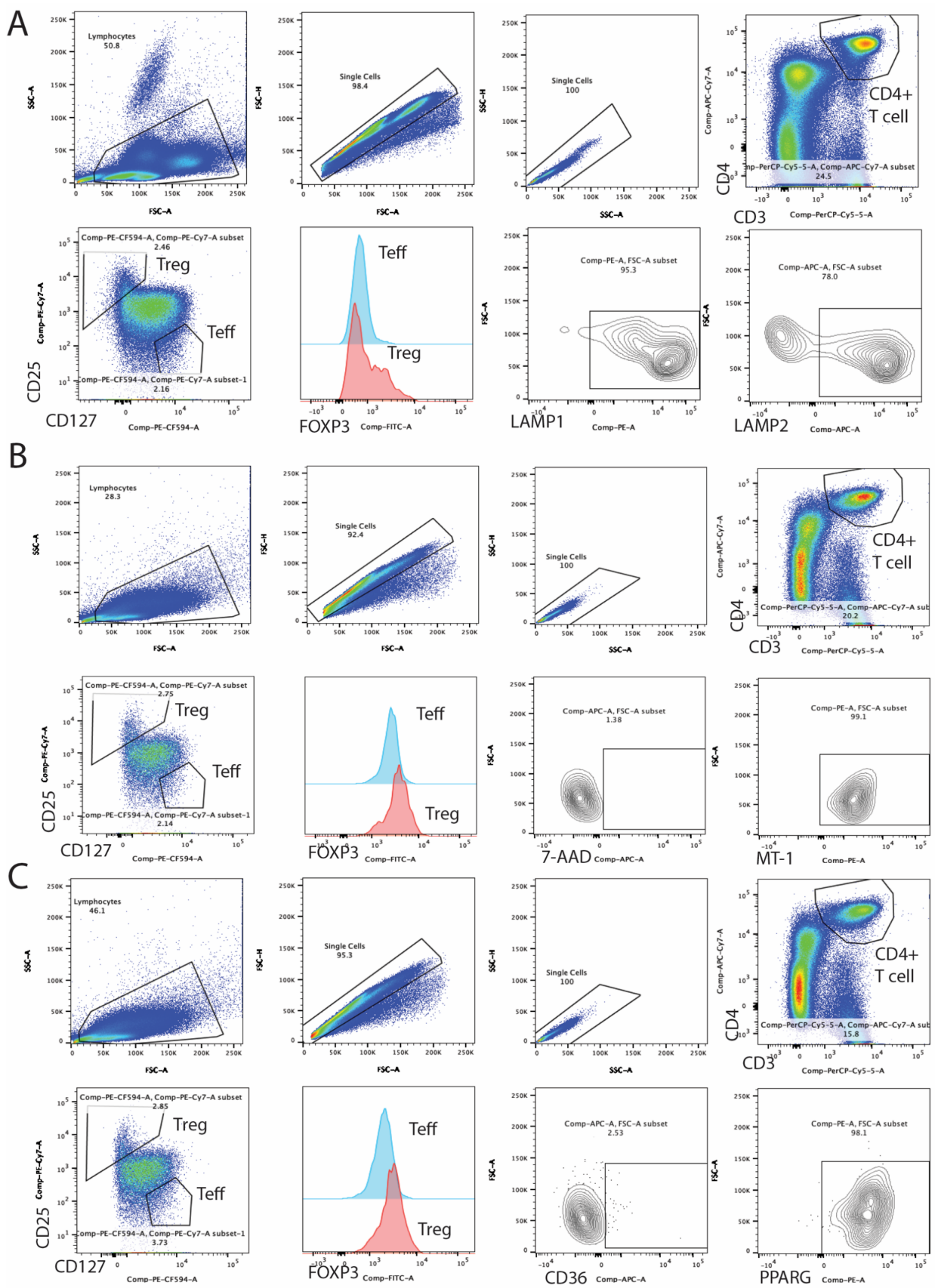


**Figure S8 Regulatory T cells have enriched lysosome, mitochondria and peroxisome.**

Regulatory T cells were analyzed with (A) lysosomal proteins (LAMP1, LAMP2) (B) mitochondrial membrane potential (MT1) (C) CD36 and PPAR-gamma (n=4).
